# Ion Mobility–Enhanced Liquid Chromatography Coupled with Mass Spectrometry (LC–MS) Enables Reliable Detection of OXA-48-Like Carbapenemases Beyond Conventional Activity-Based Assays

**DOI:** 10.64898/2026.03.30.715343

**Authors:** Vendula Studentova, Veronika Paskova, Lucia Dadovska, Jaroslav Hrabak

## Abstract

Carbapenemases are major drivers of carbapenem resistance in Gram-negative bacteria and pose a critical threat to last-line antibiotic therapy. Rapid identification of carbapenemase classes is essential for appropriate treatment and epidemiological surveillance; however, current functional methods lack class-level resolution and may yield false-negative results for OXA-48-like enzymes.

In this study, we developed and validated an assay based on liquid chromatography–mass spectrometry with trapped ion mobility spectrometry–time-of-flight [LC–MS (timsTOF)] for simultaneous detection and class-level differentiation of five clinically relevant carbapenemases (KPC, NDM, VIM, IMP, and OXA-48-like). The method employs three carbapenem substrates (meropenem, imipenem, and ertapenem).

A total of 55 clinical isolates were analyzed using a standardized 2-hour incubation protocol, with a total analysis time of 7 min per sample. Ion mobility enabled unambiguous identification of the OXA-48-specific meropenem-derived β-lactone based on its distinct collisional cross-section (185 Å² vs. 191 Å² for intact meropenem), despite identical mass and nearly identical retention time. This marker was detected in all OXA-48-like producers and was absent in all other groups. In contrast, imipenem and ertapenem did not provide comparable discrimination, highlighting the central role of meropenem. Distinct hydrolysis profiles enabled class-level differentiation supported by multivariate analysis.

LC–MS (timsTOF) thus enables rapid, sensitive, and specific functional detection of carbapenemases within a single workflow. The ion mobility dimension is critical for accurate identification of OXA-48-like enzymes and supports the potential implementation of this approach in routine clinical microbiology laboratories.

**Importance:** This study introduces an ion mobility–enabled LC–MS (timsTOF) approach for functional detection and class-level differentiation of clinically relevant carbapenemases within a single analytical workflow. By leveraging collisional cross-section measurements, the method enables reliable identification of OXA-48-like carbapenemase through detection of a meropenem-derived β-lactone that is indistinguishable by mass alone. This directly addresses a major diagnostic limitation of conventional activity-based assays, which may yield false-negative results for OXA-48-like enzymes. The approach further demonstrates the potential of integrating ion mobility into routine clinical mass spectrometry to enhance specificity beyond traditional mass and retention time measurements. These findings support the development of next-generation diagnostic strategies capable of detecting both known and emerging resistance mechanisms without reliance on predefined targets.

## Introduction

Carbapenemases are enzymes that degrade carbapenem antibiotics, which are considered last-line agents for the treatment of multidrug-resistant bacterial infections, particularly those caused by Gram-negative pathogens. Over recent decades, the global prevalence of carbapenemase-producing bacteria has increased dramatically, significantly compromising the clinical efficacy of these critical antibiotics (Bonomo et al., 2018; ECDC 2024). This resistance is predominantly associated with Gram-negative organisms, especially *Enterobacterales* (notably *Escherichia coli* and *Klebsiella pneumoniae*), as well as non-fermenting rods (e.g., *Pseudomonas* spp., *Acinetobacter* spp.).

Based on their catalytic mechanisms, most carbapenemases are classified into three molecular classes according to the Ambler classification system (A, B, and D). The major clinically relevant enzymes include KPC (class A), NDM, VIM, and IMP (class B metallo-β-lactamases), and OXA-48-like enzymes (class D). These enzymes inactivate carbapenems primarily through hydrolysis of the β-lactam ring, whereas OXA-48-like enzymes may either follow the classical hydrolytic pathway or catalyze the formation of β-lactone products, both resulting in antibiotic inactivation (Nordmann et al., 2011; Queenan et al., 2007; Studentova et al., 2022).

The increasing prevalence of carbapenemase-producing organisms is driven by multiple factors, including inappropriate antibiotic use, suboptimal infection control practices, nosocomial transmission, and global travel, all of which facilitate the dissemination of resistant strains (Nordmann et al., 2011). In response to this growing threat, new therapeutic strategies have recently been developed, including novel β-lactam/β-lactamase inhibitor combinations (e.g., avibactam, relebactam, vaborbactam) and antibiotics such as cefiderocol (McCreary et al., 2021; Petillon et al., 2025).

Accurate detection of carbapenemase-producing organisms and precise classification of carbapenemase types are essential not only for epidemiological surveillance but also for guiding targeted therapy with novel class-specific inhibitors. Currently available diagnostic approaches can be broadly divided into two groups: activity-based assays (e.g., MALDI-TOF MS hydrolysis assays, Carba NP test) and methods based on molecular or immunological detection of the gene or protein. Activity-based assays enable the detection of carbapenemase activity regardless of enzyme type, including novel variants. In contrast, molecular and immunological methods offer rapid and sensitive detection but are limited to predefined targets and cannot identify previously uncharacterized enzymes (Hrabak et al., 2014).

Direct activity-based assays may be affected by an alternative carbapenem degradation pathway—β-lactone formation—which is a distinctive feature of OXA-48-like carbapenemases observed with meropenem, ertapenem, and doripenem, but not with imipenem. Because β-lactone formation does not result in measurable changes in molecular mass or pH, such assays—particularly the Carba NP test and MALDI-TOF MS hydrolysis assays—may yield false-negative results. This limitation can be addressed by detecting β-lactone–derived products or by applying liquid chromatography–mass spectrometry (LC-MS) approaches capable of distinguishing compounds based on retention time or ion mobility (Studentova et al., 2022).

We previously demonstrated that β-lactone formation from meropenem by OXA-48-like enzymes contributes to high false-negative rates in conventional assays. Meropenem-derived β-lactone can be detected by LC–MS based on its distinct retention time (Studentova et al., 2022) and by MALDI-TOF MS as a characteristic signal (m/z 362.5), achieving high specificity in a large clinical cohort (689 isolates; sensitivity 79%, specificity 100%) (Studentova et al., 2024).

LC-MS is a well-established analytical platform widely used in clinical laboratories for therapeutic drug monitoring, toxicology, and biomarker analysis (Yu et al., 2024; Seger & Salzmann, 2020). Its increasing availability in hospital settings makes it a promising tool for the development of novel diagnostic approaches, including functional detection of carbapenemase activity.

In this study, we present a validation of an LC-MS method implemented on a Bruker timsTOF Pro 2 platform, combining high-resolution time-of-flight mass spectrometry with trapped ion mobility spectrometry [LC-MS (timsTOF)] to enable class-level differentiation of clinically relevant carbapenemases without the need for specific inhibitors. Trapped ion mobility spectrometry (TIMS) separates ions in the gas phase according to their size, shape, and charge prior to mass analysis, providing collisional cross-section (CCS) values as an additional molecular descriptor beyond retention time and accurate mass (Liu et al., 2018; Meier et al., 2018).

## Materials and Methods

### Bacterial Isolates and MIC Determination

The study included 55 fully sequenced carbapenemase-producing *Enterobacterales and Pseudomonas* isolates (Skalova et al., 2017; Chudejova et al., 2021; Kraftova et al., 2022) (Table 1). Species identification was performed using the MALDI Biotyper® system (Bruker Daltonics, Bremen, Germany). The largest group of tested carbapenemases comprised 17 isolates that produced OXA-48-like carbapenemases alone, 13 isolates co-producing OXA-48-like and NDM-type carbapenemases, reflecting the clinical relevance of this co-production phenotype. Among the remaining isolates, 9 produced NDM-type metallo-β-lactamases alone, and 6 produced KPC-type carbapenemases. In addition, five *Pseudomonas aeruginosa* isolates producing VIM and 1 producing IMP were included. As negative controls, 4 carbapenem-susceptible isolates expressing extended-spectrum β-lactamases (ESBLs) were included to verify assay specificity.

**Table 1.** Description of the isolates used in the study.

| No. | Isolate ID | Species | β-lactamase | No. | Isolate ID | Species | β-lactamase |
| --- | --- | --- | --- | --- | --- | --- | --- |
| 1 | 328 | *Escherichia coli* | KPC | 28 | 676 | *Klebsiella pneumoniae* | NDM, OXA-48 |
| 2 | 329 | *Escherichia coli* | OXA-48 | 29 | 677 | *Klebsiella pneumoniae* | NDM |
| 3 | 431 | *Pseudomonas aeruginosa* | IMP | 30 | 678 | *Klebsiella pneumoniae* | NDM, OXA-48 |
| 4 | 432 | *Escherichia coli* | KPC | 31 | 679 | *Klebsiella pneumoniae* | NDM, OXA-48 |
| 5 | 433 | *Escherichia coli* | NDM, OXA-48 | 32 | 680 | *Klebsiella pneumoniae* | NDM, OXA-48 |
| 6 | 528 | *Klebsiella pneumoniae* | KPC | 33 | 681 | *Klebsiella pneumoniae* | NDM |
| 7 | 534 | *Klebsiella pneumoniae* | NDM | 34 | 682 | *Klebsiella pneumoniae* | OXA-48 |
| 8 | 538 | *Pseudomonas aeruginosa* | VIM | 35 | 683 | *Klebsiella pneumoniae* | NDM, OXA-48 |
| 9 | 540 | *Escherichia coli* | OXA-48 | 36 | 684 | *Klebsiella pneumoniae* | OXA-48 |
| 10 | 542 | *Klebsiella pneumoniae* | NDM | 37 | 686 | *Pseudomonas aeruginosa* | VIM |
| 11 | 543 | *Klebsiella pneumoniae* | NDM, OXA-48 | 38 | 687 | *Klebsiella pneumoniae* | OXA-48 |
| 12 | 544 | *Klebsiella pneumoniae* | NDM, OXA-48 | 39 | 688 | *Klebsiella pneumoniae* | NDM, OXA-48 |
| 13 | 545 | *Klebsiella pneumoniae* | OXA-48 | 40 | 697 | *Escherichia coli* | NDM, OXA-48 |
| 14 | 546 | *Klebsiella pneumoniae* | NDM, OXA-48 | 41 | 702 | *Morganella morganii* | KPC |
| 15 | 547 | *Klebsiella pneumoniae* | NDM, OXA-48 | 42 | 703 | *Pseudomonas aeruginosa* | VIM |
| 16 | 554 | *Klebsiella pneumoniae* | KPC | 43 | 828 | *Escherichia coli* | OXA-48 |
| 17 | 601 | *Klebsiella oxytoca* | OXA-48 | 44 | 838 | *Klebsiella pneumoniae* | OXA-48 |
| 18 | 606 | *Pseudomonas aeruginosa* | VIM | 45 | 839 | *Klebsiella pneumoniae* | OXA-48 |
| 19 | 607 | *Klebsiella pneumoniae* | OXA-48 | 46 | 841 | *Enterobacter hormaechei* | OXA-48 |
| 20 | 612 | *Klebsiella pneumoniae* | OXA-48 | 47 | 842 | *Citrobacter koseri* | OXA-48 |
| 21 | 613 | *Klebsiella pneumoniae* | NDM | 48 | 843 | *Klebsiella pneumoniae* | OXA-48 |
| 22 | 631 | *Enterobacter cloacae* | NDM | 49 | 844 | *Klebsiella pneumoniae* | NDM, OXA-48 |
| 23 | 647 | *Klebsiella pneumoniae* | NDM | 50 | 845 | *Klebsiella pneumoniae* | OXA-48 |
| 24 | 648 | *Klebsiella pneumoniae* | NDM | 51 | 846 | *Klebsiella pneumoniae* | OXA-48 |
| 25 | 655 | *Pseudomonas aeruginosa* | IMP | 52 | 1833/06 | *Klebsiella pneumoniae* | SHV-2 |
| 26 | 671 | *Klebsiella pneumoniae* | NDM | 53 | 1841/06 | *Escherichia coli* | CTX-M-9 |
| 27 | 673 | *Klebsiella pneumoniae* | KPC | 54 | 1849/06 | *Enterobacter cloacae* | SHV-12, TEM-12 |
|  |  |  |  | 55 | 1850/06 | *Proteus mirabilis* | TEM-92 |

Minimum inhibitory concentrations (MICs) of ertapenem, meropenem, and imipenem were determined using broth microdilution (Erba Lachema, s.r.o., Brno, Czech Republic), and results were interpreted according to the European Committee on Antimicrobial Susceptibility Testing (EUCAST) version 15.0 criteria (www.eucast.org).

### Detection of Carbapenemases by MALDI-TOF MS

Carbapenemase activity was initially determined for all isolates using a meropenem hydrolysis assay by MALDI-TOF MS as previously described (Papagiannitsis et al., 2015). Shortly, bacterial cells were suspended in a buffer containing 20 mM Tris–HCl, 50 mM ammonium bicarbonate (NH₄HCO₃), and 0.1 mM zinc sulfate monohydrate (ZnSO₄·H₂O; Sigma-Aldrich, Prague, Czech Republic), adjusted to pH 7.0, to a density equivalent to a 3.0 McFarland standard. A 1 mL aliquot of the suspension was centrifuged, and the resulting pellet was resuspended in 50 μL of reaction buffer (20 mM Tris–HCl, pH 7.0; Sigma-Aldrich) supplemented with 0.1 mM meropenem, imipenem, or ertapenem.

Each analytical batch included a positive control (carbapenemase-producing strain with an indicator antibiotic), a negative control (carbapenemase-negative strain with an indicator antibiotic), and an antibiotic blank (antibiotic solution without bacteria) to monitor assay performance and to exclude non-enzymatic degradation. Following incubation at 35 °C for 2 h, the reaction mixture was centrifuged, and 1 µL of the supernatant was applied to a MALDI target plate and overlaid with 2,5-dihydroxybenzoic acid (10 mg/mL in 50% ethanol) as the matrix. Measurements were performed using a rapifleX MALDI-TOF mass spectrometer (Bruker Daltonics, Bremen, Germany) in positive ion mode.

### LC-MS (timsTOF) Analysis

An aliquot of 25 µL from the reaction mixture described in the previous paragraph was also subjected to LC–MS (timsTOF) analysis performed using an ultra-high-performance liquid chromatography system (Elute+ UHPLC, Bruker Daltonics, Bremen, Germany) coupled to a timsTOF Pro 2 mass spectrometer equipped with an electrospray ionization (ESI) source (Bruker Daltonics, Bremen, Germany).

Chromatographic separation was carried out on an Intensity Solo C18 reversed-phase column (100 × 2.0 mm, 2.0 µm particle size, 100 Å pore size; Bruker Daltonics, Bremen, Germany). The mobile phase consisted of 0.1% (v/v) formic acid in water (solvent A) and 0.1% (v/v) formic acid in acetonitrile (solvent B). Ultrapure water was obtained from a Milli-Q system (Millipore, USA), and LC–MS grade acetonitrile and formic acid were purchased from Sigma-Aldrich (St. Louis, MO, USA).

Gradient elution was performed as follows: 5% B for 0.3 min, increased linearly to 30% B at 1.2 min, then to 95% B at 4.0 min, followed by a hold at 95% B until 4.5 min. The column was subsequently re-equilibrated at 5% B from 4.7 to 7.0 min, resulting in a total run time of 7.0 min. The flow rate was 0.3 mL/min, the column temperature was maintained at 35 °C, and the injection volume was 5 µL. Mass spectrometric detection was performed in a positive ESI mode with full-scan acquisition over the m/z range of 20–1300. Source parameters were as follows: capillary voltage 4000 V, end plate offset 500 V, nebulizer pressure 2.2 bar, dry gas flow 10 L/min, and dry gas temperature 220 °C. For the determination of collisional cross-section (CCS) values, TIMS parameters were set to a mobility range of 1/K₀ 0.45–1.20 V·s/cm². Detailed TIMS and ion transfer parameters are provided in Supplementary Table S1.

Prior to each analytical batch, the mass spectrometer was calibrated using ESI-L Low Concentration Tuning Mix (Agilent Technologies, Santa Clara, CA, USA) to ensure mass accuracy within ±5 ppm. For each specimen, the calibrant was injected at the beginning of the analysis. Instrument control and data acquisition were performed using HyStar 6.2 and timsControl 5.0 (Bruker Daltonics, Bremen, Germany), and raw data were processed in Compass DataAnalysis 6.1. Multivariate statistical analysis, including principal component analysis (PCA), was conducted using MetaboScape 2025b (Bruker Daltonics, Bremen, Germany).

### Statistical Analysis

Multivariate statistical analysis was performed using MetaboScape (Bruker Daltonics, Bremen, Germany). Data were normalized using positive and negative control samples for batch correction and subsequently Pareto-scaled.

Principal component analysis (PCA) was used to explore unsupervised clustering patterns, while partial least squares discriminant analysis (PLS-DA) was applied to assess class separation between carbapenemase groups. Model performance was evaluated by cross-validation (CV accuracy = 68.8%, R²Y = 0.466), and variables with VIP scores >1.0 were considered discriminative.

PCA and PLS-DA were computed and visualized in Python (v3.9.6) using scikit-learn (v1.6.1), matplotlib (v3.9.4), and pandas (v2.3.3). To minimize overfitting, model robustness was assessed by cross-validation.

## Results

### Chromatographic and Mass Spectrometric Characterization of Carbapenem Hydrolysis Products

All three carbapenems (meropenem, imipenem, and ertapenem) were detected as protonated [M+H]⁺ ions in positive ESI mode, with mass accuracies within ±5 ppm of the theoretical monoisotopic masses (meropenem, m/z 384.159; imipenem, m/z 300.120; ertapenem, m/z 476.198). Representative chromatograms and corresponding MS spectra for each antibiotic and its major hydrolysis products are shown in Figure 1.

**Figure 1.**
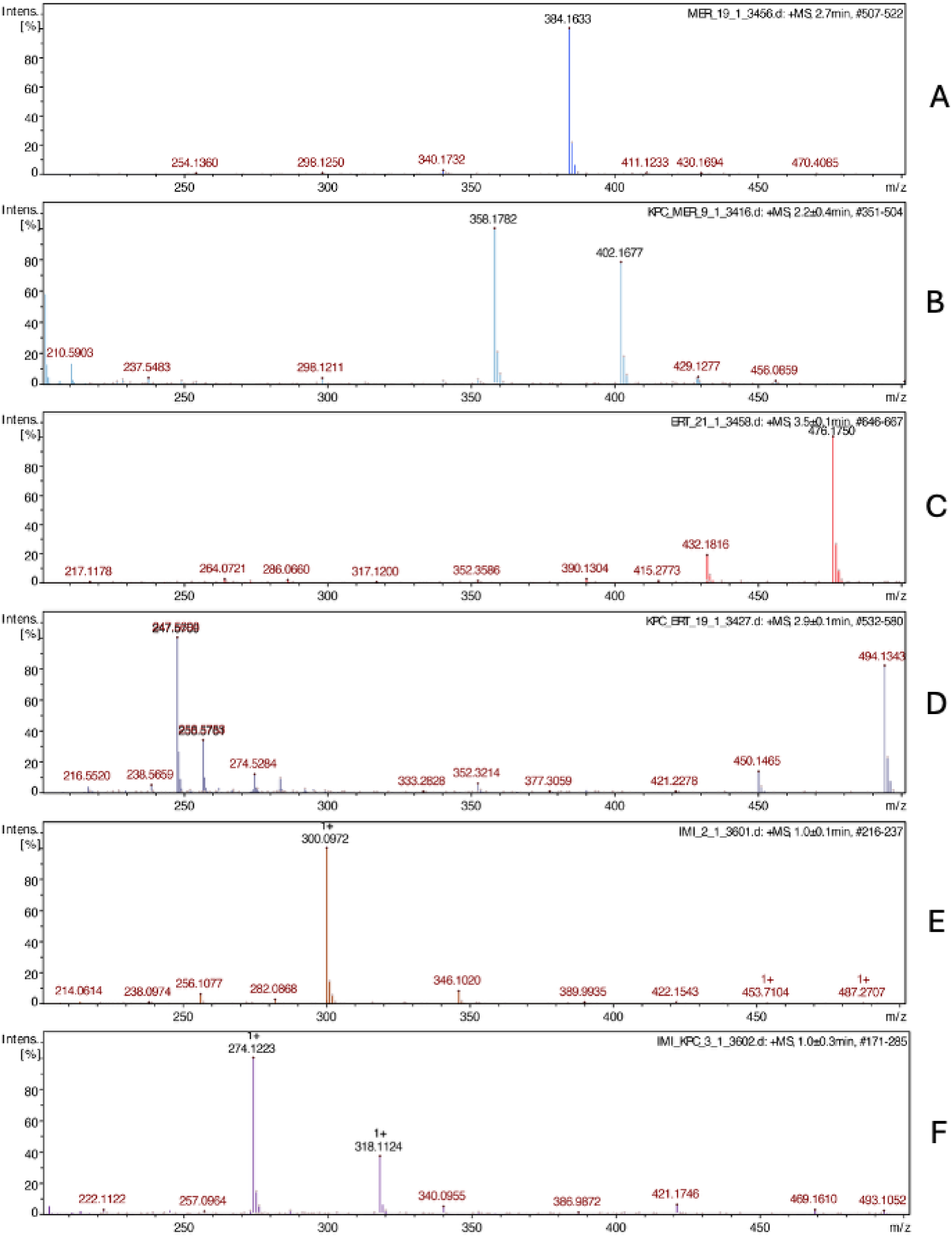
LC–MS spectra of meropenem (m/z 384) (A), its hydrolyzed (m/z 402) and decarboxylated hydrolyzed products (m/z 358) (B); ertapenem (m/z 476) (C), its hydrolyzed (m/z 494) and decarboxylated hydrolyzed products (m/z 450) (D); and imipenem (m/z 300) (E), its hydrolyzed (m/z 318) and decarboxylated hydrolyzed products (m/z 274) (F).

For class A (KPC) and class B (NDM, VIM, IMP) carbapenemases, the primary degradation pathway involved nucleophilic addition of water to the β-lactam ring, resulting in ring opening and a mass increase of +18.010 Da relative to the parent compound. For meropenem, this corresponded to a neutral mass of 401.161 Da, with analogous hydrolysis products observed for imipenem and ertapenem.

In addition, secondary products consistent with decarboxylation (−44.010 Da) were detected for meropenem and ertapenem, in agreement with previously described carbapenem degradation pathways (Studentova et al., 2022) (Figure 2).

**Figure 2.**
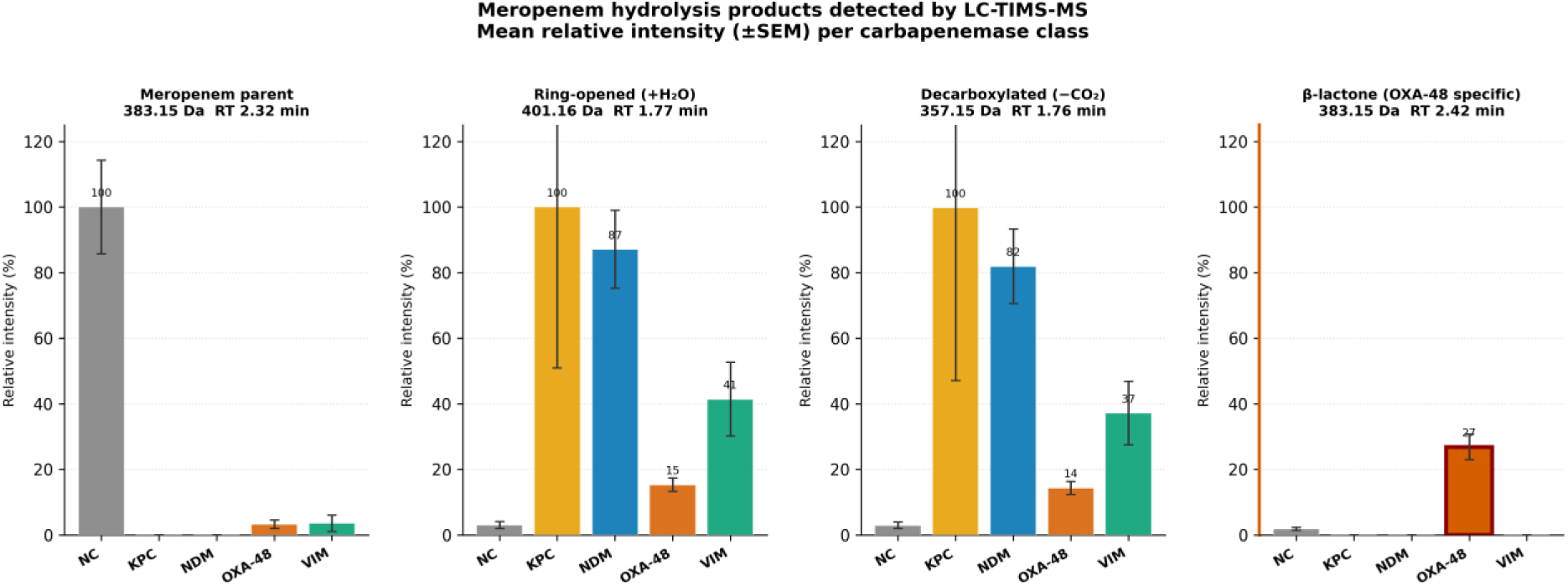
Meropenem hydrolysis products detected by LC-TIMS-MS across carbapenemase classes. Bar charts show relative abundance of intact meropenem, ring-opened hydrolysis product (+18 Da), decarboxylated product (−44 Da), and β-lactone adduct for each carbapenemase class (OXA-48-like, NDM+OXA-48 co-producers, NDM, KPC, VIM, IMP, and ESBL controls). Each panel represents one carbapenem substrate.

### Ion Mobility Separation of the OXA-48-Specific β-Lactone Product

OXA-48-like carbapenemases produced a characteristic meropenem-derived β-lactone degradation product that was nominally indistinguishable from intact meropenem by accurate mass alone (neutral monoisotopic mass 383.151 Da in both cases). Under our chromatographic conditions, the two species exhibited a retention time difference of only 0.10 min (meropenem, 2.32 min; β-lactone, 2.42 min), which may be insufficient for reliable discrimination in complex biological matrices. However, the two species were clearly resolved in the ion mobility dimension: intact meropenem displayed a collisional cross-section (CCS) of 192 Å², whereas the β-lactone product exhibited a CCS of 186 Å² (ΔCCS = 6 Å²) (Figure 3). The combined use of retention time, accurate mass, and CCS thus enabled unambiguous identification of the β-lactone.

**Figure 3.**
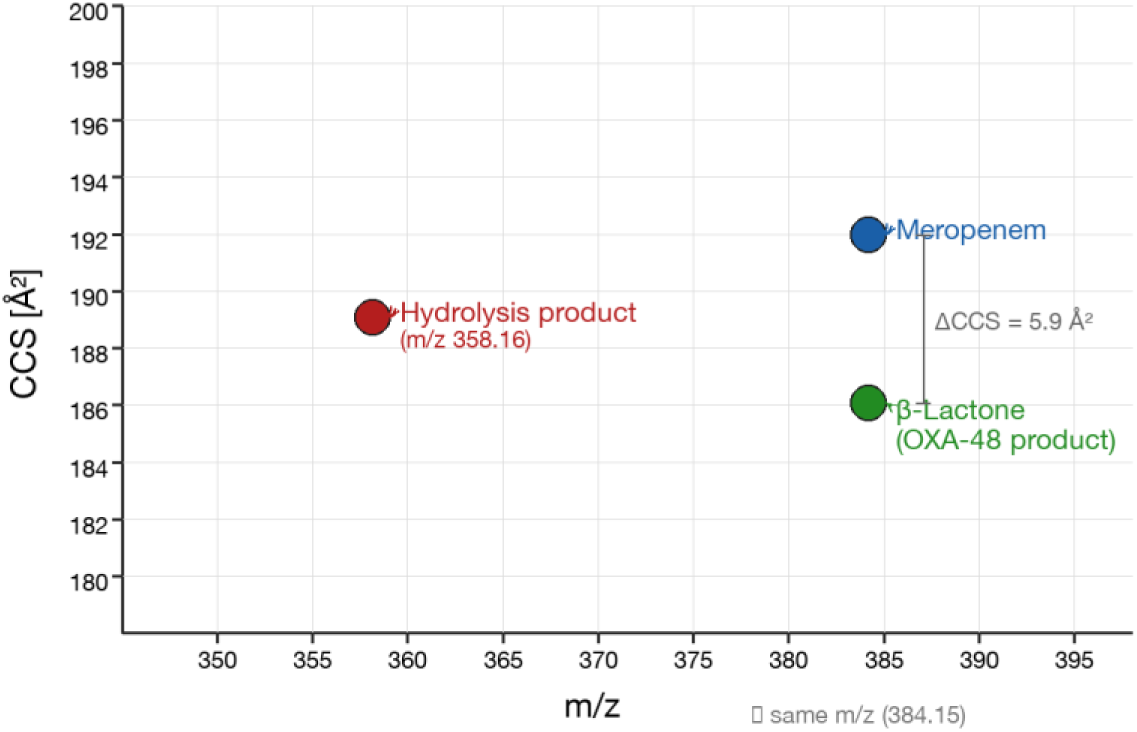
Comparison of CCS values for meropenem and its β-lactone product.

As illustrated in Figure 4, the EIC chromatogram and ion mobilogram for m/z 384.1 in the meropenem standard revealed two distinct peaks in the ion mobility dimension, corresponding to intact meropenem (1/K₀ = 0.931 V·s/cm²; CCS = 192 Å²) and the β-lactone isomer (1/K₀ = 0.901 V·s/cm²; CCS = 186 Å²). These findings confirm that, despite near-identical retention times and indistinguishable accurate masses, the two species can be reliably differentiated based on their ion mobility behavior.

**Figure 4.**
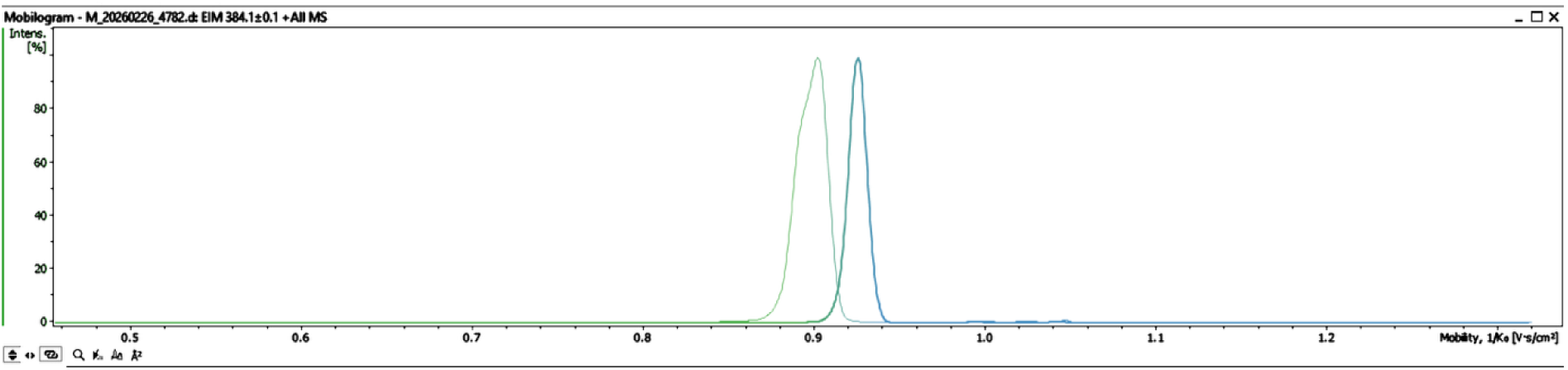
Ion mobility separation of meropenem (blue) and its β-lactone isomer (green)

By LC-MS (timsTOF), the β-lactone product was detected in all OXA-48-like-producing isolates tested, including those co-producing OXA-48-like with NDM-type enzymes, demonstrating 100% sensitivity within this cohort. It was absent from all NDM, KPC, VIM, and IMP isolates as well as from carbapenem-susceptible controls, indicating 100% analytical specificity for OXA-48-like enzymatic activity. Fourteen isolates co-producing NDM- and OXA-48-like carbapenemases were also included and clustered with OXA-48-like producers in multivariate analysis, consistent with the dominant OXA-48-like activity associated with β-lactone formation. No analogous substrate-specific β-lactone product was observed for ertapenem or imipenem, consistent with the structural requirements for β-lactone formation.

### Class-Specific Hydrolysis Profiles

All five carbapenemase classes produced distinct degradation profiles that enabled class-level differentiation. KPC-type (class A; n = 6) and NDM-type (class B; n = 9) carbapenemases caused complete hydrolysis of all three carbapenems, yielding predominantly ring-opened hydrolysis products. The resulting LC-MS (timsTOF) profiles of KPC and NDM producers were highly similar, reflecting their shared hydrolytic mechanism. Accordingly, reliable discrimination between these two groups based solely on carbapenem degradation profiles was not achieved, although partial separation was observed in multivariate PLS-DA analysis (CV accuracy = 68.8%, R²Y = 0.466) (Figure 5).

**Figure 5.**
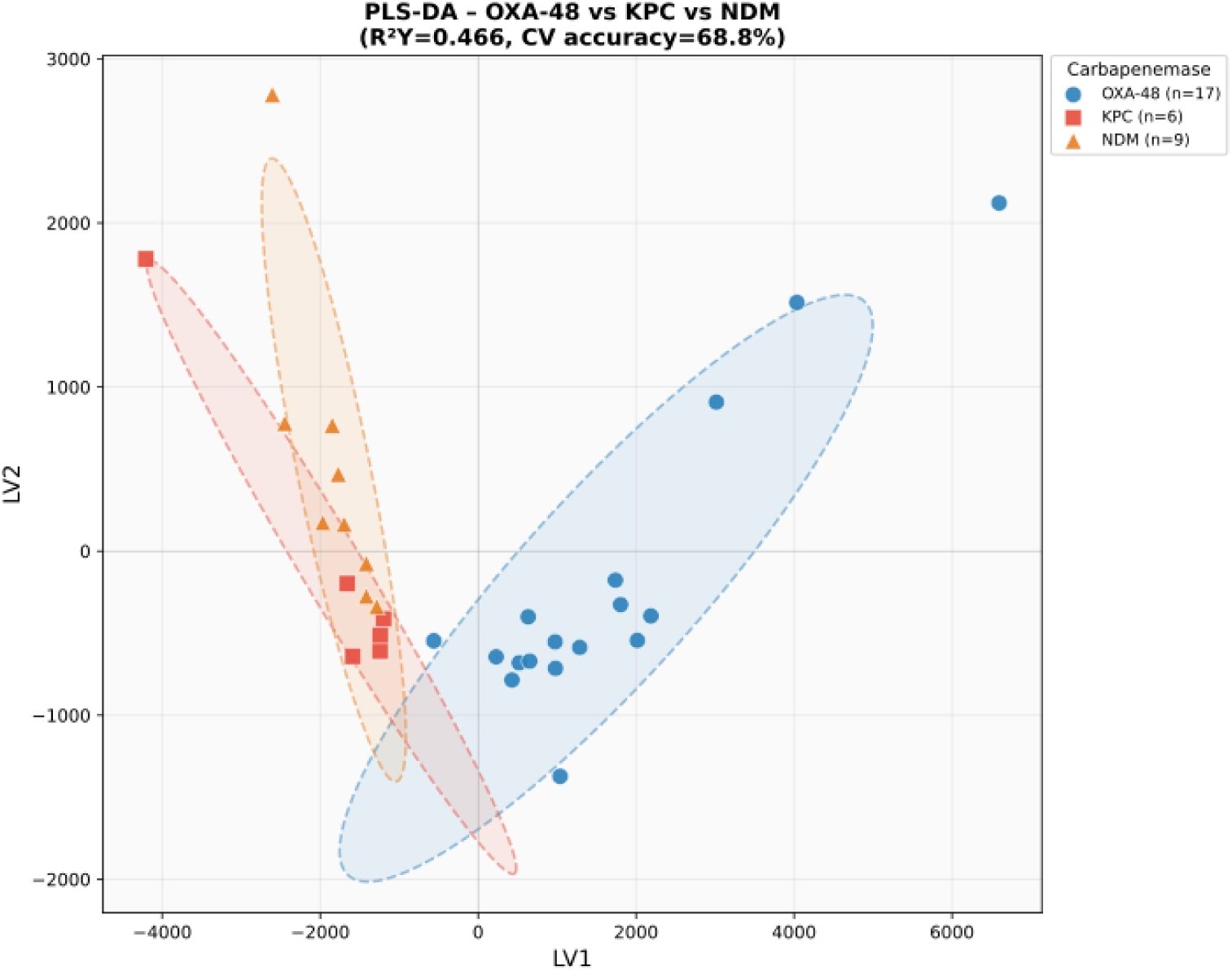
PLS-DA score plot of meropenem LC-TIMS-MS profiles for OXA-48-like (n = 17), KPC-type (n = 6), and NDM-type (n = 9) carbapenemase-producing isolates. Each point represents an individual isolate, and dashed ellipses indicate 95% confidence regions. OXA-48-like producers are clearly separated from KPC and NDM isolates along LV1, primarily driven by the presence of the β-lactone hydrolysis product (m/z 384.16, VIP = 22.3). In contrast, KPC and NDM producers overlap along both latent variables, reflecting their similar hydrolytic profiles.

VIM-type (n = 5) and IMP-type (n = 2) metallo-β-lactamases also mediated complete hydrolysis of meropenem and ertapenem, generating spectral profiles largely overlapping with those of NDM and KPC producers. Notably, all VIM-producing isolates were *Pseudomonas aeruginosa*, contributing a distinct metabolomic background that enabled their separation from *Enterobacterales* in multivariate analysis. This included organism-specific metabolites such as pyochelin and precursors of the Pseudomonas quinolone signal (PQS). Given the limited number of IMP-producing isolates (n = 2), conclusions for this group should be interpreted with caution.

Among carbapenem-susceptible control isolates carrying extended-spectrum β-lactamases (ESBLs; n = 4), variable and inconsistent imipenem degradation was observed despite the absence of carbapenemase genes, with degradation scores ranging from 0 to 0.67. In contrast, ertapenem and meropenem remained largely unaffected under the same conditions. These findings highlight the limited specificity of imipenem as a substrate in activity-based assays and support the preferential use of meropenem and ertapenem for carbapenemase detection.

### Multivariate Statistical Analysis

Principal component analysis (PCA) of the LC-MS (timsTOF) feature matrix derived from meropenem hydrolysis reactions yielded a two-component model explaining 61,2% of total variance (PC1 = 37.5%, PC2 = 23.9%; Figure 6). The PCA score plot revealed distinct spatial clustering of the major carbapenemase producer groups. OXA-48-like producers and NDM+OXA-48 co-producers formed separate, well-defined clusters, reflecting their unique β-lactone-dominated degradation profiles. KPC and NDM producers occupied overlapping regions of PCA space, consistent with the indistinguishable hydrolytic products observed at the mass spectrometric level.

**Figure 6.**
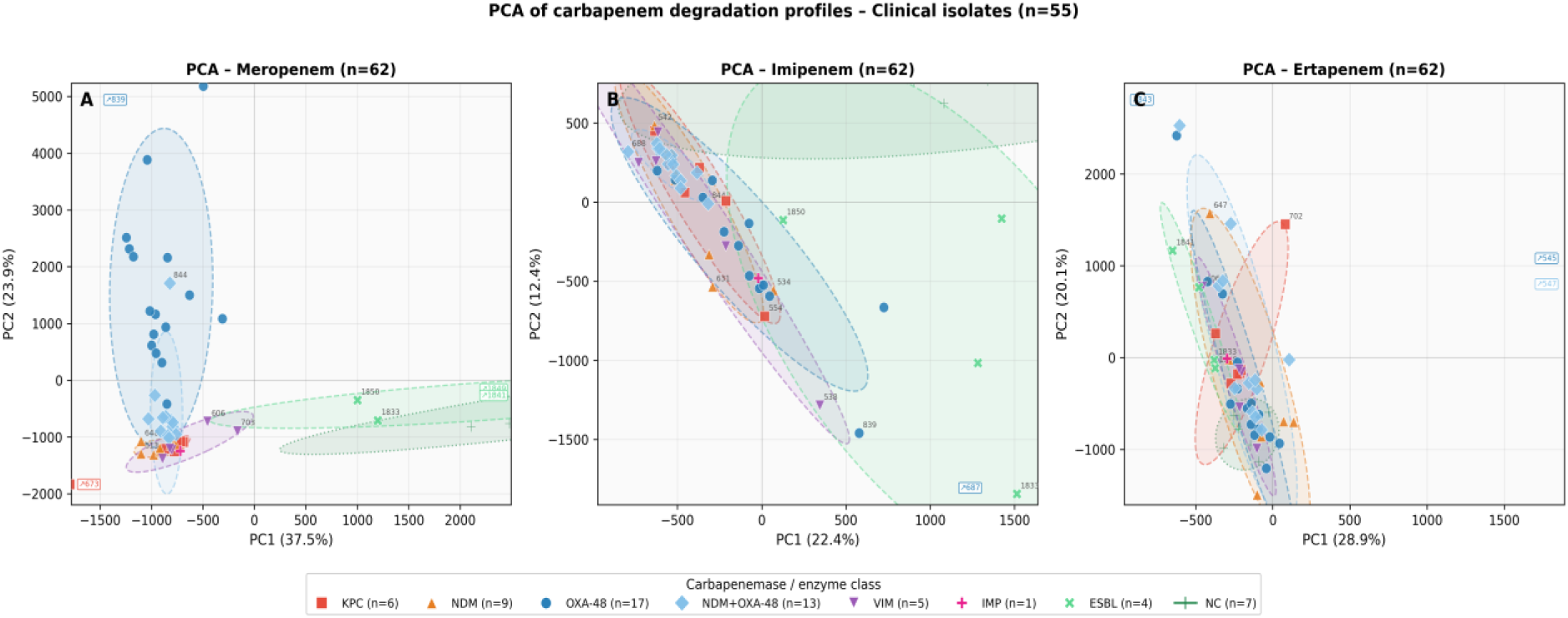
Principal component analysis (PCA) of LC-TIMS-MS feature matrices from three carbapenem substrates. Score plots show group-level discrimination across carbapenemase classes (OXA-48-like, NDM+OXA-48-like co-producers, NDM, KPC, VIM, and ESBL controls) for meropenem, imipenem, and ertapenem datasets.

VIM-producing isolates clustered distinctly from all *Enterobacterales* groups, reflecting both enzyme-specific hydrolysis activity and the divergent metabolomic background of *P. aeruginosa*. Negative control isolates (carbapenem-susceptible ESBL producers) were distributed outside the clusters of confirmed carbapenemase producers, confirming the discriminatory power of the method. Consistent results were obtained from PCA models derived from imipenem and ertapenem datasets; however, the discriminatory resolution for OXA-48-like producers was reduced in the imipenem model, reflecting the absence of β-lactone formation from this substrate.

## Discussion

OXA-48-like carbapenemases, which predominate in Europe, including the Czech Republic, remain particularly challenging to detect due to the absence of a mass shift or pH change during β-lactone formation, often leading to false-negative results in activity-based assays such as MALDI-TOF MS hydrolysis assays and the Carba NP test (Studentova et al., 2022). In this context, LC–MS (timsTOF), combining high-resolution mass spectrometry with trapped ion mobility spectrometry, enables class-level differentiation of clinically relevant carbapenemases within a single analytical workflow. The key methodological advance lies in the ion mobility-based identification of the meropenem-derived β-lactone produced specifically by OXA-48-like carbapenemases, which, although indistinguishable from intact meropenem by accurate mass, is clearly resolved by its distinct collisional cross-section (CCS; 186 vs. 192 Å²). This finding builds on previous observations (Studentova et al., 2022) and establishes a direct analytical approach for detecting OXA-48-like activity based on ion mobility characteristics rather than mass or retention time alone, thereby enabling differentiation of isobaric species based on their three-dimensional structure.

For KPC and NDM carbapenemases, hydrolysis of all three carbapenem substrates was complete, and the resulting products were indistinguishable by mass spectrometry. This reflects the convergent hydrolytic mechanisms of class A serine β-lactamases and class B metallo-β-lactamases rather than a limitation of the LC-MS (timsTOF) approach. Reliable discrimination between these classes therefore requires complementary methods, such as molecular assays or inhibitor-based phenotypic testing. The use of multiple substrates partially improved group separation in multivariate analysis, reflecting differences in substrate-specific kinetic profiles.

The substantial imipenem degradation observed in ESBL-producing control isolates highlights an important limitation of activity-based assays. Imipenem is more susceptible to non-carbapenemase-mediated hydrolysis and may therefore generate false-positive signals, particularly in strains with elevated ESBL expression. In contrast, meropenem and ertapenem exhibited higher specificity and are thus more suitable substrates for carbapenemase detection, consistent with previous reports (Papagiannitsis et al., 2015).

VIM-producing isolates in this study were exclusively *Pseudomonas aeruginosa*, and their separation in PCA space was influenced by both enzyme activity and the organism-specific metabolomic background, including metabolites such as pyochelin and PQS-related quinolones. While this facilitates discrimination from Enterobacterales, it also indicates that part of the separation is taxonomically driven. Validation in VIM-producing Enterobacterales will be required to assess the enzyme-specific contribution independently of host metabolomics.

Following the initial description of a MALDI-TOF MS-based approach for carbapenemase detection (Hrabak et al., 2011), several studies demonstrated that LC–MS can also detect carbapenemase activity through analysis of indicator carbapenems and their hydrolyzed products (Carvalhaes et al., 2013; Peaper et al., 2013). However, this approach offers limited advantage over the widely available and cost-effective MALDI-TOF MS.

In contrast, LC–MS (timsTOF) introduces additional analytical dimensions that support its potential clinical implementation. The short analysis time (7 min per sample) is compatible with high-throughput workflows, and the combined evaluation of retention time, accurate mass, and CCS provides a robust multi-dimensional identification framework. Importantly, the method does not rely on prior knowledge of carbapenemase variant, enabling detection of novel or atypical enzymes that may escape genotypic or immunological assays.

With the increasing availability of LC-MS instrumentation in clinical laboratories, particularly in clinical biochemistry and therapeutic drug monitoring settings, implementation of LC-MS-based diagnostic assays in microbiology is becoming increasingly feasible. Although the method requires access to high-resolution instrumentation with ion mobility capability, sample preparation is straightforward and does not require specialized reagents beyond standard carbapenem substrates. This study represents a proof of concept demonstrating the potential of LC–MS (timsTOF) for the functional differentiation of carbapenemases. While the results are encouraging, further validation in larger, multicenter cohorts will be necessary to confirm robustness across diverse clinical settings. Discrimination between certain carbapenemase classes remained less pronounced, suggesting that complementary approaches may be beneficial in specific contexts. Notably, the present work primarily focuses on OXA-48-like carbapenemases, for which the method provides a clear analytical advantage. Future studies will therefore aim to expand the range of evaluated enzymes and to assess the applicability of this approach in clinically relevant workflows.

## Acknowledgement

The study was supported by Charles University Grant Agency (GA UK), project Nr. 280323, by the project „Integration of biomedical research and health care in the Pilsen metropolitan area“; reg. no. CZ.02.01.01/00/23_021/0008828) - co-funded by the European Union and by the State Budget of the Czech Republic, and by the project Nr. CZ.02.1.01/0.0/0.0/16_019/0000787 “Fighting Infectious Diseases” provided by the Ministry of Education Youth and Sports of the Czech Republic.

## References

Nordmann, P., Naas, T., & Poirel, L. (2011). Global spread of Carbapenemase-producing Enterobacteriaceae. Emerging infectious diseases, 17(10), 1791–1798. 10.3201/eid1710.110655

Bonomo, R. A., Burd, E. M., Conly, J., Limbago, B. M., Poirel, L., Segre, J. A., & Westblade, L. F. (2018). Carbapenemase-Producing Organisms: A Global Scourge. Clinical infectious diseases : an official publication of the Infectious Diseases Society of America, 66(8), 1290–1297. 10.1093/cid/cix893

European Centre for Disease Prevention and Control. 2025. Antimicrobial resistance in the EU/EEA (EARS-Net): annual epidemiological report 2024. ECDC, Stockholm.

Queenan, A. M., & Bush, K. (2007). Carbapenemases: the versatile beta-lactamases. Clinical microbiology reviews, 20(3), 440–458. 10.1128/CMR.00001-07

Studentova, V., Sudova, V., Bitar, I., Paskova, V., Moravec, J., Pompach, P., Volny, M., Novak, P., & Hrabak, J. (2022). Preferred β-lactone synthesis can explain high rate of false-negative results in the detection of OXA-48-like carbapenemases. Scientific reports, 12(1), 22235. 10.1038/s41598-022-26735-5

Petillon, C., Ulutuipalelei, M., Beyrouthy, R., Proust, S., Aissa, N., Bret, L., Brieu, N., Carricajo, A., Cattoir, V., Dauwalder, O., Degand, N., Dortet, L., Doucet-Populaire, F., Dubois, V., Grillon, A., Guérin, F., Lanotte, P., Lemaitre, N., Leyssene, D., Neuwirth, C., … Robin, F. (2025). Evaluation of imipenem-relebactam, meropenem-vaborbactam, aztreonam-avibactam and cefepime-zidebactam activities on a wide collection of French clinical Enterobacterales isolates. Diagnostic microbiology and infectious disease, 113(3), 117011. 10.1016/j.diagmicrobio.2025.117011

McCreary, E. K., Heil, E. L., & Tamma, P. D. (2021). New Perspectives on Antimicrobial Agents: Cefiderocol. Antimicrobial agents and chemotherapy, 65(8), e0217120. 10.1128/AAC.02171-20

Hrabák, J., Chudáčková, E., & Papagiannitsis, C. C. (2014). Detection of carbapenemases in Enterobacteriaceae: a challenge for diagnostic microbiological laboratories. Clinical microbiology and infection : the official publication of the European Society of Clinical Microbiology and Infectious Diseases, 20(9), 839–853. 10.1111/1469-0691.12678

Studentova, V., Dadovska, L., & Hrabak, J. (2024). Direct identification of OXA-48-type carbapenemases by detection of β-lactone-specific signal using matrix-assisted laser desorption/ionization time-of-flight (MALDI-TOF) mass spectrometry. International journal of antimicrobial agents, 63(5), 107130. 10.1016/j.ijantimicag.2024.107130

Yu, S., Zou, Y., Ma, X., Wang, D., Luo, W., Tang, Y., Mu, D., Zhang, R., Cheng, X., & Qiu, L. (2024). Evolution of LC-MS/MS in clinical laboratories. Clinica chimica acta; international journal of clinical chemistry, 555, 117797. 10.1016/j.cca.2024.117797

Seger, C., & Salzmann, L. (2020). After another decade: LC-MS/MS became routine in clinical diagnostics. Clinical biochemistry, 82, 2–11. 10.1016/j.clinbiochem.2020.03.004

Liu, F. C., Ridgeway, M. E., Park, M. A., & Bleiholder, C., (2018). Tandem trapped ion mobility spectrometry. The Analyst, 143(10), 2249–2258. 10.1039/c7an02054f

Meier, F., Brunner, A. D., Koch, S., Koch, H., Lubeck, M., Krause, M., Goedecke, N., Decker, J., Kosinski, T., Park, M. A., Bache, N., Hoerning, O., Cox, J., Räther, O., & Mann, M. (2018). Online Parallel Accumulation-Serial Fragmentation (PASEF) with a Novel Trapped Ion Mobility Mass Spectrometer. Molecular & cellular proteomics: MCP, 17(12), 2534–2545. 10.1074/mcp.TIR118.000900

Skalova, A., Chudejova, K., Rotova, V., Medvecky, M., Studentova, V., Chudackova, E., Lavicka, P., Bergerova, T., Jakubu, V., Zemlickova, H., Papagiannitsis, C. C., & Hrabak, J. (2017). Molecular Characterization of OXA-48-Like-Producing Enterobacteriaceae in the Czech Republic and Evidence for Horizontal Transfer of pOXA-48-Like Plasmids. Antimicrobial agents and chemotherapy, 61(2), e01889–16. 10.1128/AAC.01889-16

Chudejova, K., Kraftova, L., Mattioni Marchetti, V., Hrabak, J., Papagiannitsis, C. C., & Bitar, I. (2021). Genetic Plurality of OXA/NDM-Encoding Features Characterized From Enterobacterales Recovered From Czech Hospitals. Frontiers in microbiology, 12, 641415. 10.3389/fmicb.2021.641415

Kraftova, L., Finianos, M., Studentova, V., Chudejova, K., Jakubu, V., Zemlickova, H., Papagiannitsis, C. C., Bitar, I., & Hrabak, J. (2021). Evidence of an epidemic spread of KPC-producing Enterobacterales in Czech hospitals. Scientific reports, 11(1), 15732. 10.1038/s41598-021-95285-z

Papagiannitsis, C. C., Študentová, V., Izdebski, R., Oikonomou, O., Pfeifer, Y., Petinaki, E., & Hrabák, J. (2015). Matrix-assisted laser desorption ionization-time of flight mass spectrometry meropenem hydrolysis assay with NH4HCO3, a reliable tool for direct detection of carbapenemase activity. Journal of clinical microbiology, 53(5), 1731–1735. 10.1128/JCM.03094-14

Hrabák, J., Walková, R., Studentová, V., Chudácková, E., & Bergerová, T. (2011). Carbapenemase activity detection by matrix-assisted laser desorption ionization-time of flight mass spectrometry. Journal of clinical microbiology, 49(9), 3222–3227. 10.1128/JCM.00984-11

Carvalhaes, C. G., Cayô, R., Assis, D. M., Martins, E. R., Juliano, L., Juliano, M. A., & Gales, A. C. (2013). Detection of SPM-1-producing Pseudomonas aeruginosa and class D β-lactamase-producing Acinetobacter baumannii isolates by use of liquid chromatography-mass spectrometry and matrix-assisted laser desorption ionization-time of flight mass spectrometry. Journal of clinical microbiology, 51(1), 287–290. 10.1128/JCM.02365-12

Peaper, D. R., Kulkarni, M. V., Tichy, A. N., Jarvis, M., Murray, T. S., & Hodsdon, M. E. (2013). Rapid detection of carbapenemase activity through monitoring ertapenem hydrolysis in Enterobacteriaceae with LC-MS/MS. Bioanalysis, 5(2), 147–157. 10.4155/bio.12.310

